# Continuum Theory for Planar Cell Polarity

**DOI:** 10.1101/2021.11.30.468758

**Authors:** Mohd Suhail Rizvi, Divyoj Singh, Mohit K. Jolly

## Abstract

Planar Cell Polarity (PCP), characterized by asymmetric localization of proteins at the cell membrane within the epithelial plane, plays essential roles in embryonic development and physiological functions. The significance of PCP can be appreciated by the outcomes of its failure in the form of defects in neural tube formation, tracheal malfunctions, organ shape misregulation, hair follicle misalignment etc. Extensive experimental works on PCP in fruit fly *Drosophila melanogaster* have classified the proteins involved in PCP into a ‘core’ module, acting locally by inter-cellular protein interactions, and a ‘global’ module, responsible for the alignment of cell polarities with that of the tissue axis. Despite the involvement of different molecular players the asymmetric localization of the proteins of the two modules on cell membrane primarily involves inter-cellular dimer formations. We have developed a continuum model of the localization of PCP proteins on the cell membrane and its regulation via intra- and inter-cellular protein-protein interactions. We have identified the conditions for the asymmetric protein localization, or PCP establishment, for uniform and graded protein expression levels in the tissue. We have found that in the absence of any tissue level expression gradient polarized state of the tissue does not arise. However, in the presence of tissue-level expression gradients of proteins the polarized state remains stable. We have also looked at the influence of the loss of PCP proteins from a select region of the tissue on the polarization of the cells outside of that region. This continuum theory of planar cell polarity can be coupled with active-matter hydrodynamics to study cell flows and their regulation by genetic machinery.

## 1 Introduction

### 1.1 Biological context

In epithelial tissues, which form the outer surface of most organs, cells are polarized in two directions-apico-basal [1, 2], which is the direction normal to the epithelial surface, and within the plane of the epithelium, also known as planar cell polarity (PCP)[3, 4, 5]. The mechanisms of the establishment of PCP have been investigated extensively using experimental (see [5] for Review) and theoretical approaches (see [6] for Review). Experimental investigations, primarily performed using the fruit fly *Drosophila melanogaster* as the model organism, have revealed the asymmetric distribution of several proteins/protein-complexes at the apical periphery of the epithelial cells to be the primary mechanism of PCP [5, 7, 8]. The importance of the planar cell polarity in epithelial tissues can be appreciated by the fallout of the PCP disruptions which encompass several developmental abnormalities including defects in neural tube formation [9, 10, 11], ciliopathies [12, 13], tracheal ciliary malfunctions [14], misregulation of organ shape maintenance [15], hair follicle misalignment [16].

#### 1.1.1 Proteins involved in the planar cell polarity

The main molecular players involved in PCP have been characterized and divided into two sets [7, 4], known as ‘global’ and ‘core’ (or local) modules of PCP. The global module, also known as Ft-Ds-Fj pathway, involves two atypical cadherins Fat (Ft) and Dachsous (Ds), a kinase Four-jointed (Fj), and myosin Dachs (D) [15]. Among these, Ft and Ds, both present in phosphorylated and unphosphorylated forms, from two neighboring cells have mutual affinity and form inter-cellular heterodimers at the cell-cell interface [17, 5]. Fj modulates the Ft-Ds affinity by their phosphorylation: phosphorylation of Ft increases it while that of Ds decreases it [5, 4]. The expression pattern of Ft, Ds and Fj is regulated by morphogen gradients in the tissue and as a result they too display respective tissue-level expression gradients [18]. The final member of the module Dachs co-localizes with Ds and regulates the organs shape by generating active forces at the apical end [15].

On the other hand, the ‘core’ module of the PCP consists of Frizzled (Fz), Flamingo (Fmi, also known as Starry night), Van gogh (Vang), Prickle-spiny-legs (Pk), Dishevelled (Dsh) and Diego (Dgo) [19] among which Fmi, an adhesion-GPCR transmembrane cadherin, plays the central role in inter-cellular communication [20, 21]. Given the complexity of the module the exact nature of the interaction among its member proteins is not completely understood [22]. However, as per the current understanding, Fmi from two neighboring cells form homo-dimers at the cell-cell interface (akin to the Ft-Ds heterodimer in the global module) [7, 5, 23]. The other core proteins provide functional asymmetry to two Fmi molecules forming the dimer by interacting with them in two different ways [5, 22]. The transmembrane protein Fz recruits Dsh and Dgo (both cytosolic proteins) and localizes close to Fmi in one of the two cells [5, 23]. Vang, another transmembrane proteins, recruits cytosolic protein Pk to localize close to the Fmi on the other cell [7, 5, 23]. There is also evidence [24] that Vang and Fz in two cells interact directly without the intermediary Fmi.

#### 1.1.2 Mechanisms of PCP establishment

Despite the diversity in the players involved, the primary mechanisms of the establishment of asymmetry in protein localization in both of these modules have been shown to have similarities where they involve feedback system between the two proteins(in the global module)/protein-complexes(in the core module) forming the dimer at the two cell boundaries [25, 18] along with dynamic trafficking of proteins [14] and their differential localization on the membrane [26, 27]. The involvement of the inter-cellular protein dimers in the PCP establishment is one of the necessary components since PCP needs to be coordinated at the tissue scale.

In the case of the core module, the two Fmi molecules forming the inter-cellular Fmi-Fmi dimers recruit Fz or Vang at the cell membrane. This recruitment of Fz or Vang gives a polarity to the otherwise Fmi homodimer. It has also been seen that a Fmi-Fz complex in one cell is more likely to form a inter-cellular dimer with Fmi alone or Fmi-Vang complex of the neighboring cell, and not Fmi-Fz complex [28, 26]. The possible mechanism of preferential binding results in an functional asymmetry in Fmi which is propagated across the tissue. This asymmetry is further amplified by Dsh, Dgo and Pk, the remaining members of the core module, as follows. Dsh and Dgo are recruited to the Fmi-Fz complex at the cell membrane and promote further recruitment of their own species and restrict the binding of Vang and Pk to the respective Fmi along with providing junctional stability of the Fmi-Fz-Dsh-Dgo complex [29]. Similarly, Pk binds to the Fmi-Vang complex and promotes further enrichment of its own type (more Fmi-Vang-Pk) and inhibits the binding of the complex of opposite polarity (that is Fmi-Vang-Dgo-Dsh) [30]. Therefore, in addition to the functional asymmetry in the protein molecules forming the inter-cellular dimers (the inter-cellular or non-autonomous mechanism) the core module also has positive and negative feedbacks (the intra-cellular or cell-autonomous mechanism) which further amplify the asymmetric localization of the proteins. These cell-autonomous and non-autonomous mechanisms, although sufficient for the asymmetric localization of the proteins of the core module, cannot propagate PCP to very large distance and also fail to coordinate with the tissue axis. The coordination of the PCP direction with the tissue axis is mediated by the global module or Ft-Ds-Fj pathway [18].

As mentioned earlier, like the core module, the global module also acts via the formation of Ft-Ds dimers between two neighboring cells. The coordination of the PCP with the tissue axis by the global module has been shown to be regulated by the tissue-level expression gradient of the members of the global module. In several tissues Ds and Fj have been seen to be expressed in opposing gradients [31]. The tissue level expression gradients act as the global cue to the PCP molecules. The coordination between the functions of the two modules has been the least explored aspect of PCP. Still, there have been some works to show the interaction of the two modules by the differential localization of core module components with that of the global module via to isoforms of Pk-Prickle (Pk) and Spiny legs (Sple). These two isoforms interpret the Ft and Ds localization in the opposing manner [32, 33]. It has been found that the Pk isoform prefers to get localized with Ft resulting in colocalization of Fmi-Vang-Pk with Ft on the cell membrane. On the other hand, the Sple isoform prefers Ds and D resulting in colocalization of Fmi-Vang-Sple with Ds. There is also evidence to that the two modules act independently of each other since perturbation in the both the modules together shows higher disruption to the PCP as compared to that when any one of the two modules is perturbed [34]. These two possible mechanisms of interaction between core and global modules suggest either ‘series’ and ‘parallel’ action of the two modules and the debate on this aspect of PCP is not yet settled [35, 34].

### 1.2 Mathematical modeling of PCP and the current state of the art

Most of the theoretical approaches for the study of PCP have worked in a discrete cellular framework which is analyzed by computational means [6]. In one of the earliest works, focusing on the dynamics of core module, it was shown that it is the cell-autonomous negative feedback in the form of Pk blocking the recruitment of Dsh by Fz to the cell membrane which can explain some of the experimental observations including the non-autonomous effects [36]. In another theoretical work on the core module, it was proposed that the feedback is not between the proteins but the dimers formed at the cell edges such that the dimer of one polarity (say, Fmi-Fz:Fmi:Vang) promotes further formation of its own type while also inhibiting the formation of the dimer of opposite polarity (Fmi-Fz:Fmi:Vang) [37]. The same heterodimer-heterodimer feedback mechanism was further shown to explain the gradient-sensing mechanism in the global module [38]. In a phenomenological model the establishment of the planar cell polarity and its regulation by the global cue was studied using an approach motivated by models of ferromagnetism and no molecular details were taken into account [39]. Several other computational works have also looked into different aspects of the cell-autonomous and non-autonomous protein-protein interactions and have invoked feedback mechanisms to explain a range of experimental observations [40, 41].

Although such computational approaches do help in unraveling the molecular interactions central to the PCP establishment and explain the experimental observations, they do not provide any analytical insight into the PCP mechanism. This makes the comparison between PCP and other well studied physical systems, such as magnetism [37, 39], only qualitative. In this work, we propose a general continuum theory for the planar cell polarity where details of molecular interactions are taken into account. We focus on the common mechanism of inter-cellular dimer formation which is deployed by both of the PCP modules, core as well as global. We take into account cell-autonomous as well as non-autonomous protein interactions into account to obtain the necessary and sufficient conditions on the establishment of PCP. The aim of this work is to present a generic continuum framework with minimal free parameters which can be extended to specific instance of PCP by incorporating details of protein-protein interactions and cell mechanics.

## 2 Continuum model for PCP

### 2.1 Basic assumptions

#### 2.1.1 Tissue and cell geometry

We consider the epithelial tissue to be a continuum (one or two-dimensional) made of a confluent monolayer of cells with identical apical cross-sectional geometry (Fig. 1D). This coarse-grained treatment of the epithelium, as opposed to the extensively studied discrete cellular arrangement, is one of the major departures of the current work from the previously studied theoretical models of planar cell polarity [36, 37, 38, 40]. Towards our goal of a minimal model to capture PCP establishment driven by protein-protein interaction, we also assume the cells in the tissue to be non-motile and of identical geometries. Apart from very few works (where cell packing was a factor of interest) [42, 43, 44], the perfect hexagonal packing and static nature of the cells have been common assumptions across all discrete models of the PCP studies [36]. Our continuum description of the tissue describes spatial features of PCP on length scales much larger than typical diameter *l* of the apical cross-section of a cell.

**Figure 1:**
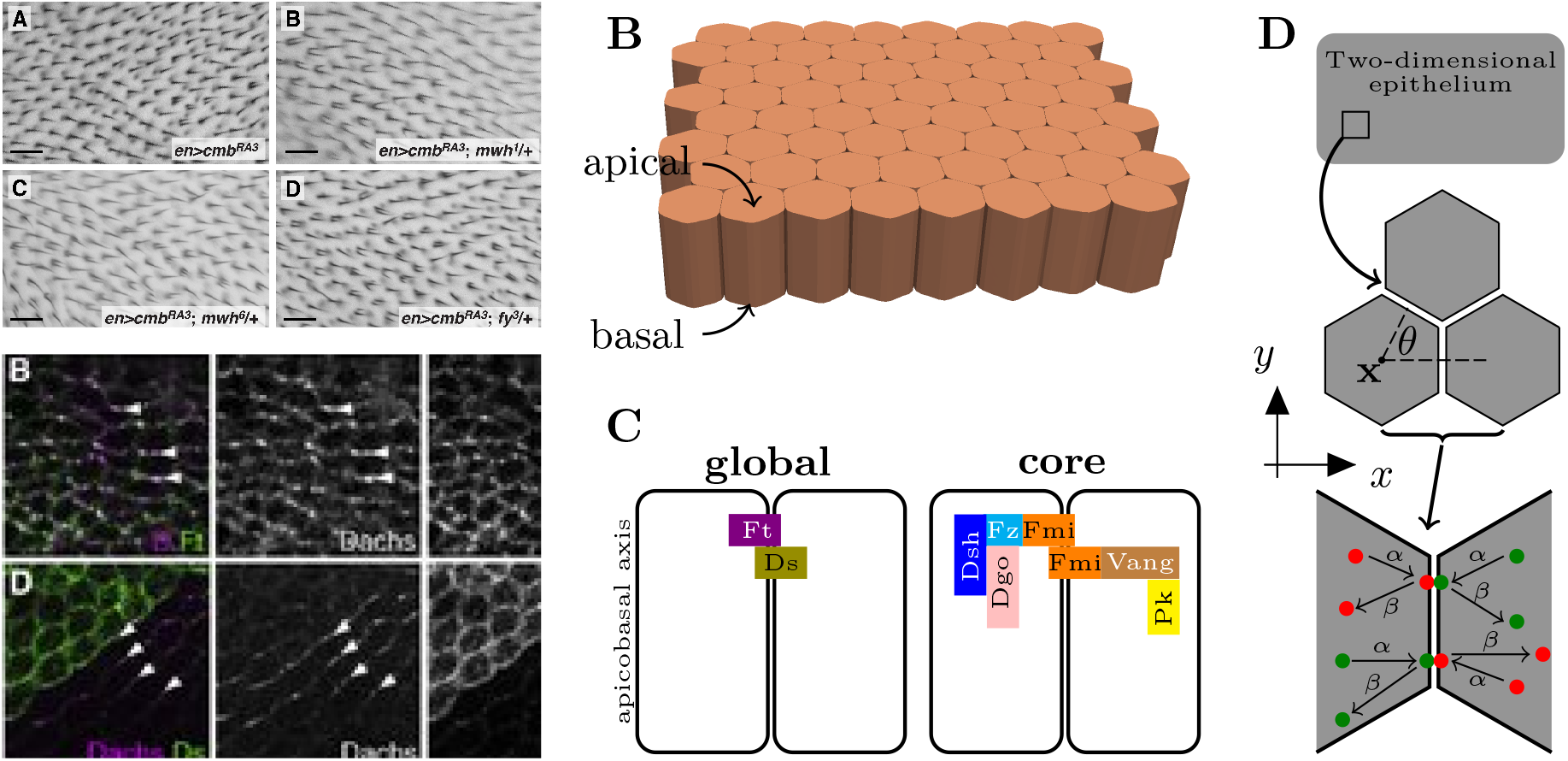
Planar cell polarity in epithelium. (A) Orientation of hairs on *Drosophila* wing epithelium (upper panel) is determined by the asymmetric localization (lower panel) of the proteins of planar cell polarity pathway. Schematics showing (B) three dimensional structure of columnar epithelium and (C) inter-cellular interactions among proteins of global and core modules of planar cell polarity pathway. (D) Two dimensional continuum representation of epithelial tissue.

#### 2.1.2 Planar cell polarity proteins

As described earlier, we consider the establishment of PCP in the form of asymmetric distribution of the PCP proteins on the cell membranes (Fig. 1C). For simplicity, we consider a system of two proteins (or protein complexes), say *a* and *b*, which can interact with each other on the membrane of a single cell (intracellular interaction) or across the cell boundaries of two cells (inter-cellular interactions). In the context of the global module of the PCP, these two proteins can be considered to be atypical cadherins Ft and Ds (Fig. 1C), and in case of the core module they represent Fmi-Vang and Fmi-Fz protein complexes (Fig. 1C). The extension of the present approach to incorporate more details of molecular interactions among proteins, for example recruitment of Dsh, Pk, Vang etc. in core module and Fj driven phosphorylation of Ft and Ds in global module, is not too difficult.

#### 2.1.3 Protein binding and transport kinetics

We assume that *a* (similarly *b* as well) present in the cytoplasm can get attached to the cell membrane with rate *α* (Fig. 1D). Similarly, *a* already attached to the cell membrane can get detached to the cytoplasm with a constant rate *β*. Here we have taken same rates *α* and *β* for both proteins but it can be easily generalized to different rates as well. We further assume that the time scale of the protein translation and degradation is much larger than that of the binding and unbinding kinetics of the two proteins and, therefore, we do not consider any temporal changes in the total protein quantities in the cells. Similarly, the transport of these proteins in the cytoplasm is taken to be purely diffusive in nature with a characteristic timescale much smaller than that of the protein binding kinetics. As a result, the protein levels are considered to be uniform within a cell and we do not take into account any intra-cellular protein gradients.

There have been some experimental evidence of active transport of core PCP components via aligned microtubules [14, 45]. This mode of active transport, however, has been shown to be regulated by the global module [46]. In the present work we do not take this active transport into account.

Note that in order to keep the continuum model minimal we are also not considering any intra-cellular interaction (the autonomous mechanisms in case of core module) such as the one presented in [47] and [38]. However, the model can be easily extended to incorporate those interactions as well.

#### 2.1.4 Inter-cellular protein interactions

Inspired by the experimental observations of the interaction between two proteins at the interface of two adjacent cells, we consider inter-cellular interaction between *a* of one cell with *b* of neighboring cell and vice versa. The effect of this inter-cellular interaction is taken into account by the reduction in the detachment rate, *β*, in the following manner. For an isolated cell the detachment of protein *a* from cell membrane to cytoplasm is *β* which, in the presence of interaction of *a* with *b* of neighboring cell, reduces to *β* (1 − *f* (*b*_*n*_)) where *f* (*b*_*n*_) is a function (for example, Hill’s function) of the amount of membrane-bound *b* in the neighboring cell at the location shared by this cell and satisfies 0 ≤ *f* (*b*_*n*_) ≤ 1. In the same manner, the reduction in the detachment rate of *b* due to its binding with *a* is also taken into account.

It needs to be pointed out that we are not interested in the actual levels of inter-cellular *a* − *b* dimers but take its effect into account in the form of reduced detachment rates of the two proteins. This is in contrast to the earlier theoretical models which focused on the dimer levels and their polarity (*a*-*b* vs *b*-*a* dimers) formed by the two proteins/protein-complexes between the adjacent cells as the primary ingredient of the establishment of PCP [40, 38]. The deviation from an analysis based on dimer levels is motivated by two factors. First, the focus on the protein distribution along cell membrane gives us much simpler system from analytical as well as mechanistic point of view. Second, even though the dimer formation has been shown conclusively there is not yet, in our knowledge, any experimental evidence to suggest that it is the dimer itself which affects the downstream PCP proteins. Further, in a recent work on the global PCP module it has been shown that the levels of Ft on the cell membrane are stabilized by the presence of Ds in the adjacent cell and vice versa [48]. In other words, inter-cellular dimer works as a mechanism (a sufficient condition) for stabilization of the PCP proteins at cell membranes, and not as a necessary ingredient for downstream PCP functioning. Experimental testing of this claim will require a setup where proteins can get localized asymmetrically without formation of heterodimer which is not easy at the present moment. Before we present the two-dimensional continuum framework for PCP it is worthwhile to consider the much simpler one-dimensional case (continuum counterpart to numerous discrete models of PCP [40, 41]).

### 2.2 One-dimensional model

In a one-dimensional tissue, the cells are described by their two edges (left *L* and right *R*). We consider the inter-cellular protein interactions to be described by

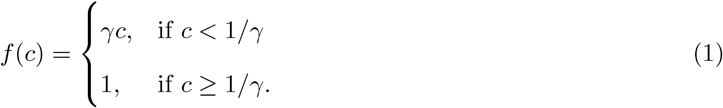

where *c* is the protein quantity at the membrane of neighboring cell. This piece-wise linear nature of *f* (*c*) is taken to simplify analytical calculations. We describe the membrane bound quantities of protein by *a*_*L*_(*x*) (amount of *a* on left edge of cell), *a*_*R*_(*x*), *b*_*L*_(*x*) and *b*_*R*_(*x*), where *x* is the cell coordinate. We utilize following quantities for non-dimensionalization of variables

1. *τ* = 1*/β* for time,
2. *l* for lengths, and
3. 1*/γ* for protein levels.

We write equations for *a*_0_(*x*) = *a*_*L*_(*x*) +*a*_*R*_(*x*) and *b*_0_(*x*) = *b*_*L*_(*x*) +*b*_*R*_(*x*), the total membrane bound proteins in a cell at location *x*, as

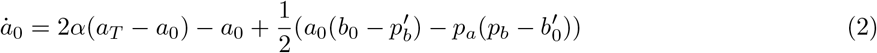

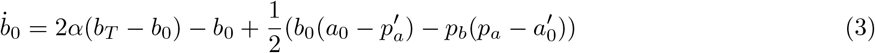

where dot (·) and prime (′) represent the derivatives with respect to non-dimensionalized time and space variables, respectively. The terms *a*_*T*_ and *b*_*T*_ stand for the amount of total (cytoplasmic+membrane bound) proteins in the cell and can be dependent on *x*, the cell location in the tissue. We have defined the asymmetries or polarities *p*_*a*_ = *a*_*R*_ − *a*_*L*_ and *p*_*b*_ = *b*_*R*_ − *b*_*L*_ in the localization of each species of protein on cell edges, a measure of PCP. In order to ensure that the protein levels do not become negative at any of the cell edges, this definition of protein asymmetry requires |*p*_*a*_| ≤ *a*_0_ and |*p*_*b*_| ≤ *b*_0_. The equations for *p*_*a*_ and *p*_*b*_ can be written as

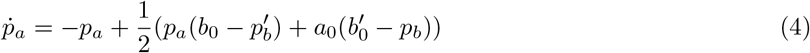

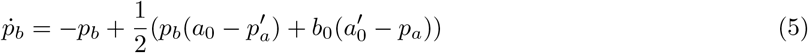

subjected to the constraints |*p*_*a*_| ≤ *a*_0_ and |*p*_*b*_| ≤ *b*_0_.

#### 2.2.1 Uniform protein expressions

For the uniform expressions of proteins *a* and *b* in the tissue, that is *a*_*T*_ (*x*) = *b*_*T*_ (*x*) = *ρ* (a known constant value), we seek homogeneous solutions. For this state, taking a difference of equations (2) and (3) and setting all spatial derivatives to zero gives us *a*_0_ = *b*_0_ = *ρ*_0_ (an unknown) for the steady state, and we obtain

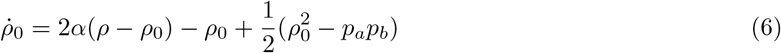

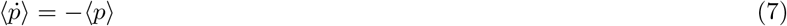

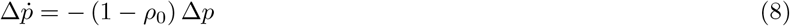

where ⟨*p*⟩ = *p*_*a*_ + *p*_*b*_ and Δ*p* = *p*_*a*_ − *p*_*b*_. These equations show that ⟨*p*⟩ = 0 at the steady state, implying localization of two proteins in opposite directions. In terms of the magnitude of the polarity the tissue can attain two states-unpolarized (Δ*p* = 0) for *ρ*_0_ ≤ 1, and polarized (Δ*p* > 0) for *ρ*_0_ > 1. In the absence of the two constraints, the last equation shows that for *ρ*_0_ > 1 the magnitude of the cell polarity increases indefinitely. However, due to |*p*_*a*_| ≤ *a*_0_ and |*p*_*b*_| ≤ *b*_0_ it can be seen that for *ρ*_0_ > 1 the two proteins get localized on the two opposite edges of the cells.

#### 2.2.2 Non-uniform protein expressions

In almost all of the epithelial tissues with PCP the members of global module show a tissue level expression gradients [49, 18, 50]. In order to study their effect we consider a weak linear [50] gradient of the expression of *a* such that *a*_*T*_ = *ρ* + *ϵx* (with *ϵ* ≪ *ρ*). We assume the following ansatz *a*_0_ = *ρ*_0_ + *ax, b*_0_ = *ρ*_0_ + *bx* with homogeneous polarization level in the tissue. For this case, we get

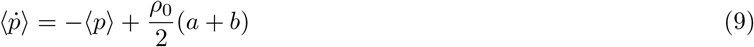

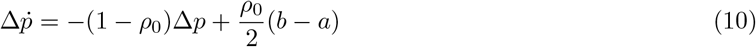

where

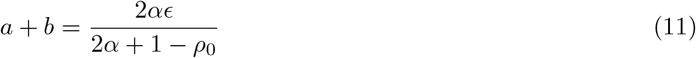

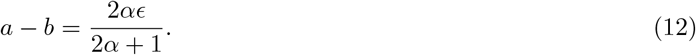

For the steady state, we obtain for *ρ*_0_ ≪ 1

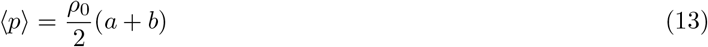

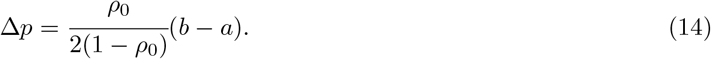

Thus a tissue-level gradient expression of one of the two proteins has an effect formally identical to that of an aligning field in magnetism: the tissue acquires a nonzero polarization in a direction determined by that of the gradient.

#### 2.2.3 Loss of one protein from a region of the tissue

In the analysis so far we have considered an infinite 1D tissue. In order to simulate the experimental condition of loss of *a* (or *b*) from a select group of cells in otherwise wildtype tissue with uniform expressions of *a* and *b*, we consider a semi-infinite tissue (that is *x* ≥ 0) where *x* = 0 corresponds to the boundary of the region where *a* is not expressed. The loss of *a* in the region *x* < 0 will set the boundary conditions for the unknown variables *a*_0_, *b*_0_, *p*_*a*_ and *p*_*b*_. In this case, we expect *a*_0_, *b*_0_, *p*_*a*_ and *p*_*b*_ to be *x*-dependent and their derivatives with respect to *x* cannot be ignored. Still, we should get the wildtype behavior as seen above for *x* ≫ 1. We assume solutions of type *a*_0_(*x*) = *ρ*_0_ + *a*(*x*), *b*_0_(*x*) = *ρ*_0_ + *b*(*x*), *p*_*a*_(*x*) = *p* + *p*_*a*_(*x*) and *p*_*b*_(*x*) = −*p* + *p*_*b*_(*x*). For keep the analysis simpler we also assume |*p*| ≪ *ρ*_0_. Substituting these values in the equations (2) and (3) gives us for the steady state

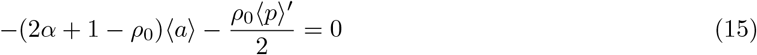

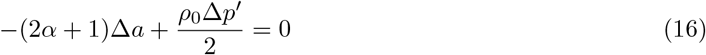

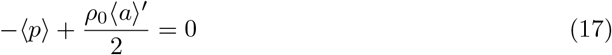

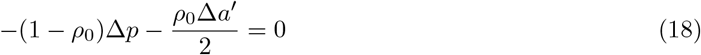

where ⟨*p*⟩ = *p*_*a*_ + *p*_*b*_, ⟨*a*⟩ = *a* + *b*, Δ*p* = *p*_*a*_ − *p*_*b*_ and Δ*a* = *a* − *b*. From these equation we obtain

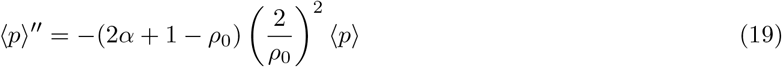

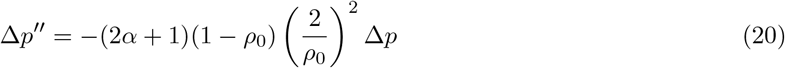

In order to ensure ⟨*p*⟩ → 0 and Δ*p* → 0 as *x* → ∞, we need *ρ*_0_ > 1 which gives

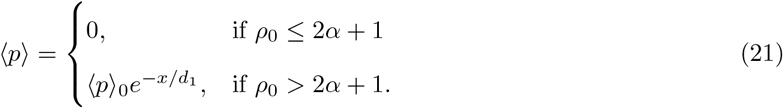

and

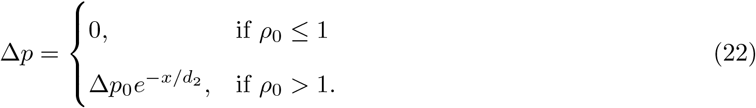

where 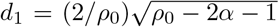 and 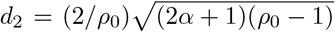, and ⟨*p*⟩_0_ and Δ*p*_0_ can be determined by the boundary condition at *x* = 0. This demonstrates an exponential decay in the magnitude of cell polarization from the boundary of the region where one of the two proteins is not expressed. For more detailed numerical analysis of this one dimensional model reader is referred to [51].

It needs to be pointed out, however, that despite capturing some of the characteristics of PCP, including loss/gain of PCP proteins from a region of the tissue, there are some experimental observations, such as non-uniform swirling orientations of the PCP in cells [5, 52, 18] or the effect of cell packing on PCP [42], which are not possible to be captured by the one dimensional framework. For this, we turn to the two dimensional continuum model in the following.

### 2.3 Two dimensional model

For simplicity, we consider the epithelium to be planar (in general epithelia can be of curved geometries and complex topology) two-dimensional surface where location of any cell is denoted by **x** (Fig. 1D). Further, we parameterize any location on the cell boundary by polar angle *θ* (Fig. 1D). Therefore, *a* (**x**, *θ, t*) denotes the distribution of the protein *a* along the periphery of the cell located at **x** in the tissue at time *t*. With this notation we can write the equations for the binding kinetics of the two proteins on the cell membrane as

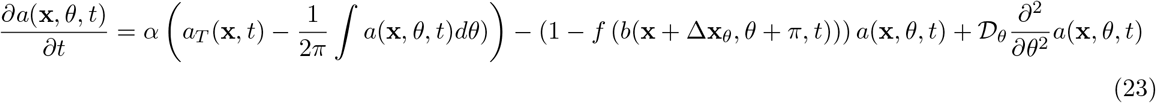

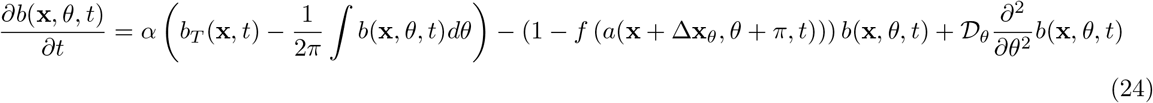

where *a*_*T*_ (**x**, *t*) and *b*_*T*_ (**x**, *t*) are the total (cytoplasmic+membrane bound) protein levels in the cell located at **x**, and *a*(**x** + Δ**x**_*θ*_, *θ* + *π, t*) represents the protein *a* in the neighboring cell (located at **x** + Δ**x**_*θ*_ with Δ**x**_*θ*_ = *l* (cos *θ*, sin *θ*)^⊤^) bound at location opposing to *θ*. In the following we consider *f* (*c*) to be same as that in the one dimensional model given by equation (1). The last term in both of the above equations correspond to the protein diffusion on the cell membrane with 𝒟_*θ*_ being the coefficient of diffusion. Since the cell boundary is periodic in *θ* we can express the protein distribution along the cell boundary in the form of Fourier series

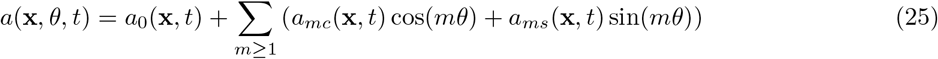

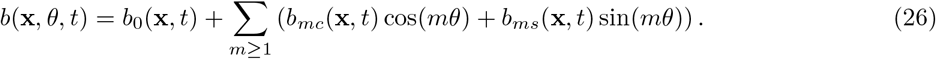

We can define the planar cell polarity (PCP) of a protein in a cell in terms of the asymmetric distribution of that protein on the cell membrane. This definition of PCP for protein *a* translates as a vector with following two components

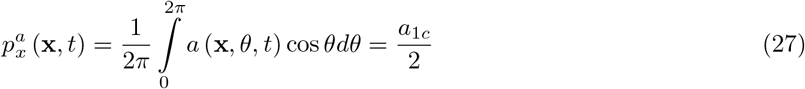

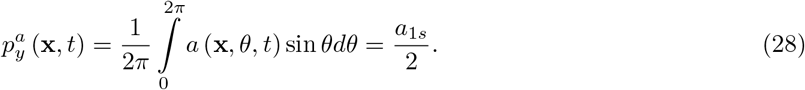

This shows that the asymmetry in the protein localization is characterized by the first harmonics of the Fourier series expansion of the protein distribution on the membrane. We substitute the Fourier series representations of *a*(**x**, *θ, t*) and *b*(**x**, *θ, t*) into equations (23) and (24) and consider upto second harmonics to write the evolution equations for *a*_0_, *b*_0_, *a*_*ic*_, *a*_*is*_, *b*_*ic*_ and *b*_*is*_ for *i* = 1, 2. By assuming slow evolution of the second harmonic we can perform adiabatic elimination to obtain

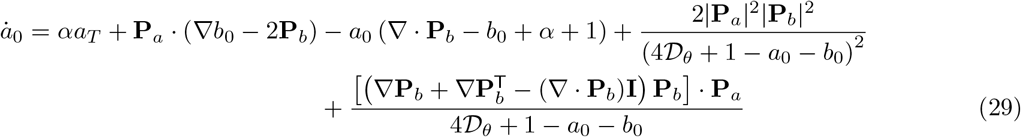

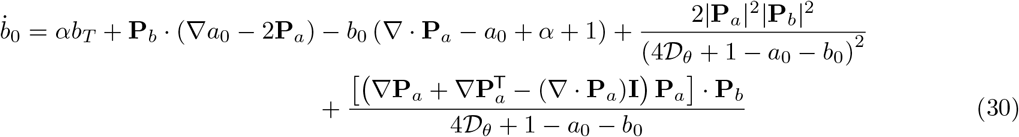

and

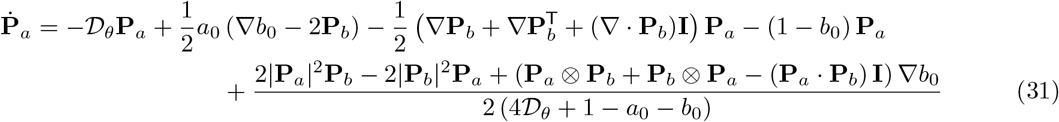

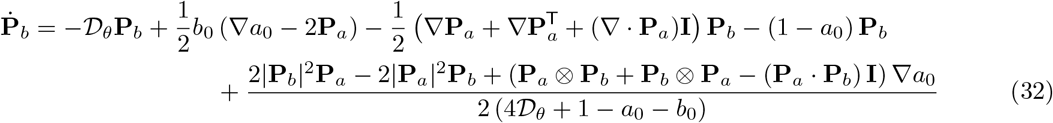

where **P**_*a*_ is the cell polarization vector with two components given by equations (27) and (28) for protein *a* (similarly **P**_*b*_ for protein *b*), and ⊗ stands for the dyadic product of two vectors. It is noticeable that this set of partial differential equations is not solvable analytically. Therefore, in the following, we will look at some special scenarios for which these equations can be approximately solved analytically.

## 3 Results

### 3.1 Uniform expression of the two proteins

#### 3.1.1 Homogeneous planar cell polarization

##### 0 Equal expression levels of two proteins

We first look at simpler case when the levels of both proteins in all the cells are same and also equal to each other, that is *a*_*T*_ = *b*_*T*_ = *ρ* (where *ρ* is some known constant value). For these expression levels we seek the homogeneous solutions to the system of equations derived above. For homogeneous solution, we can set all spatial derivative terms in the above system of equations to zero. Therefore, in the steady state we obtain from equations (29) and (30)

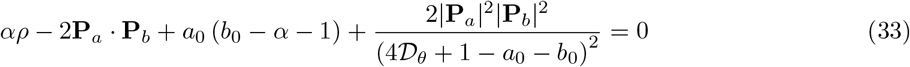

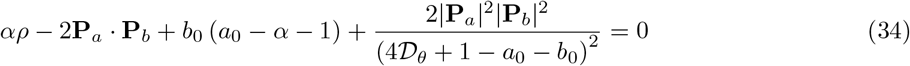

Taking the difference of these equations gives us *a*_0_ = *b*_0_ = *ρ*_0_ (where *ρ*_0_ is yet an unknown). Defining ⟨**P**⟩ = **P**_*a*_ + **P**_*b*_, Δ**P** = **P**_*a*_ − **P**_*b*_ and adding equations (31) and (32) gives us

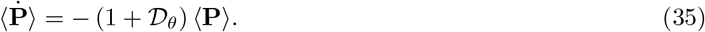

This shows that as *t* → ∞, ⟨**P**⟩ → 0, implying opposing polarization of *a* and *b*. Similarly, taking a difference of equations (31) and (32) results in

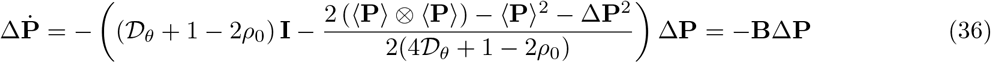

where ⊗ denotes the dyadic product and **I** is the identity tensor. By utilizing that fact that ⟨**P**⟩ = **P**_*a*_+**P**_*b*_ = 0 at the steady state, we find two fixed points of the system (also see Fig. 2A)

**Figure 2:**
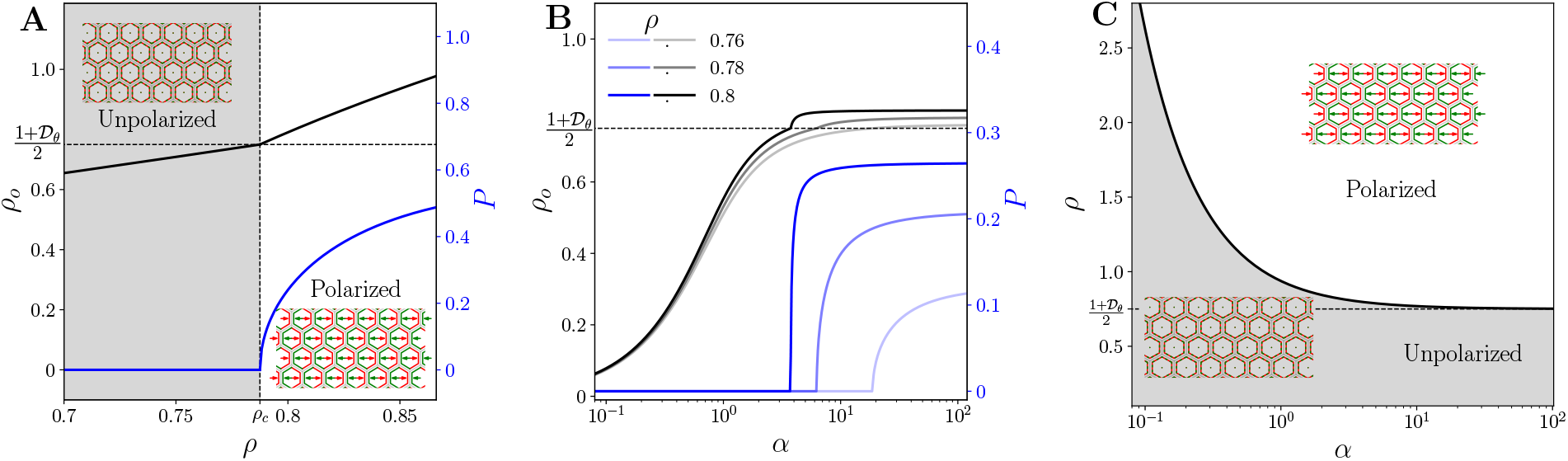
Homogeneous planar cell polarization with uniform protein expressions. (A) *ρ*_0_ and PCP magnitude *P* as a function of *a*_*T*_ = *b*_*T*_ = *ρ* for a tissue where proteins *a* and *b* are expressed equally in each cell. For *ρ* < *ρ*_*c*_, the system remains in unpolarized state. Here *α* = 5.0 and 𝒟_*θ*_ = 0.5. (B) Dependence of *ρ*_0_ and *P* on *α*, the protein binding rate to the membrane. (C) Phase diagram showing polarized and unpolarized states of the epithelial tissue as a function of total protein concentrations *ρ* and binding rate *α*.

1. Unpolarized state: ⟨**P**⟩ = 0 and |Δ**P**| = 0 for 2*ρ*_0_ < 1 + 𝒟_*θ*_
2. Polarized state: ⟨**P**⟩ = 0 and Δ**P**^2^ = 2 (4𝒟_*θ*_ + 1 − 2*ρ*_0_) (2*ρ*_0_ − 1 − 𝒟_*θ*_) for 2*ρ*_0_ > 1 + 𝒟_*θ*_

It can be seen that the condition on Δ**P** in the homogeneously polarized state only restricts its magnitude and not the direction. That means in the absence of any ‘global cue’ the tissue can have a uniformly polarized state but the polarization direction is arbitrary.

Since in both steady states we have ⟨**P**⟩ = 0, we set **P**_*a*_ = −**P**_*b*_ = **P** (where **P** is still an unknown with |**P**| = *P*). Substituting these steady state values in equation (33) or (34) gives

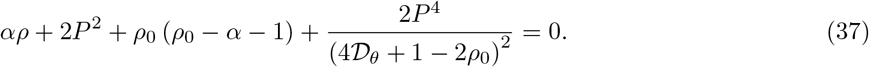

We need to solve the above equation for *ρ*_0_ along with

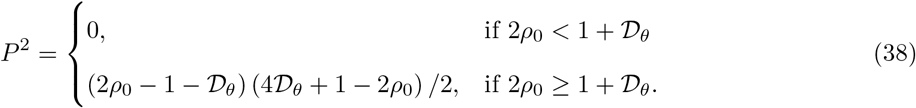

To solve this first we set *P* = 0 in the equation (37) and obtain

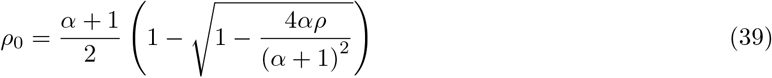

which satisfies 2*ρ*_0_ < 1 + 𝒟_*θ*_ for

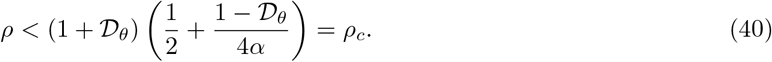

This *ρ*_*c*_ gives the critical value of total protein levels in the cell required for the polarity establishment. Fig. 2B shows the dependence of *ρ*_0_ and magnitude of cell polarity as a function of the binding rate *α*. This shows that the critical protein levels in the cells decrease with the increasing *α*. This is not surprising since an increase in the binding rate of the protein results in higher protein concentrations, which in turn, result in cell polarization at lower values of *ρ*, the total protein availability in the cell. It needs to be highlighted here that in this model we have only three free parameters among which *α* and *ρ* can be altered in the experiments with relative ease as compared to 𝒟_*θ*_. Therefore, we also plotted the phase diagram of the PCP on *ρ*-*α* plane (Fig. 2C). This shows that higher values of both *α* and *ρ* can lead to cell polarization.

In order to understand the nature of bifurcation at *ρ* = *ρ*_*c*_ we estimate the values of *ρ*_0_ and polarization magnitude *P* in the vicinity of *ρ*_*c*_, that is for *ρ* = *ρ*_*c*_ + *δρ*. We substitute 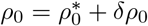 in the equation (37) where 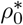 is the total bound protein for *ρ* = *ρ*_*c*_. For this we obtain

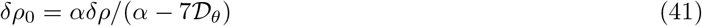

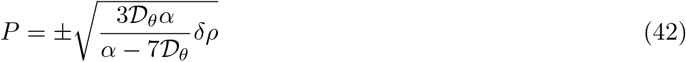

This shows the 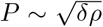 onset for the polarization magnitude, which signifies that the spontaneous polarization of the two proteins at *ρ* = *ρ*_*c*_ happens via super-critical pitchfork bifurcation.

### Stability Analysis

We also performed linear stability analysis to test if the homogeneously polarized and unpolarized states are stable against homogeneous and non-homogeneous perturbations. It can be seen from Eqs. (35) and (36) that the steady state solutions are stable against homogeneous perturbations. The stability of the homogeneous solutions, however, needs to be checked against non-homogeneous perturbations with finite length scale. We consider perturbations of type

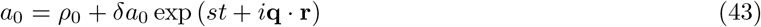

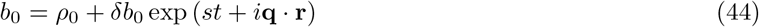

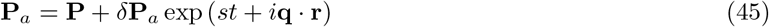

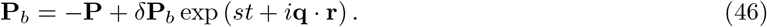

We substitute these values in the equations (29), (30), (31) and (32) and linearize these equations. After solving the linearized equations for *δa*_0_, *δb*_0_, *δ***P**_*a*_ and *δ***P**_*b*_ we obtain the dispersion relation to see the nature of perturbed solutions. For unpolarized state, that is *P* = 0, and the in the polarized state very close to the critical point *ρ*_*c*_, the dispersion relation (after ignoring quadratic terms in *P, q*) is

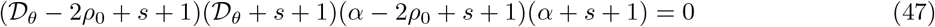

which has two negative roots (*s* = −1 − *α* and *s* = −1 − 𝒟_*θ*_) and two roots (*s* = −1 − 𝒟_*θ*_ + 2*ρ*_0_ and *s* = −1 − *α* + 2*ρ*_0_) which can be positive or negative depending on *ρ*_0_. We can see that for the unpolarized state (which satisfies 2*ρ*_0_ < 1 + 𝒟_*θ*_) all roots are negative. However, for the polarized state we get *s* = 2*ρ*_0_ − 1 − 𝒟_*θ*_ > 0 indicating the unstable nature of the polarized solution close to the critical point. We also confirmed the stability of the polarized state by numerical simulation of the equations (23)-(24) on a discrete version of this model (See [51] for more details) in the presence of noise. Noise resulted in disrupting the coordinated tissue polarization confirming the analytical results. This outcome is similar to the behavior of the polar active matter where homogeneous polarization remains unstable against non-homogeneous perturbations [53, 54]. However, in the active matter the homogeneous unpolarized state state is also unstable which is not the case here.

### Unequal expression levels of two proteins

We will now look at the scenario when the two proteins are expressed uniformly in the tissue but their levels are disparate, that is *a*_*T*_ ≠ *b*_*T*_. For homogeneous solutions in this case, after following similar steps as in previous section, we obtain

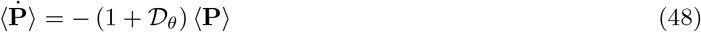

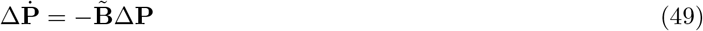

where ⟨**P**⟩ = **P**_*a*_ + **P**_*b*_, Δ**P** = **P**_*a*_ − **P**_*b*_, and

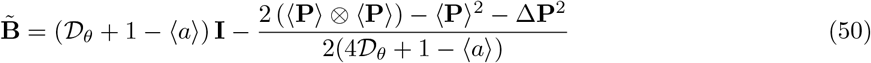

where ⟨*a*⟩ = *a*_0_ + *b*_0_. This shows that for disparate but uniform protein levels also, the polarization of two proteins in opposite in direction but equal in magnitude in each cell at steady state. By utilizing the fact that ⟨**P**⟩ = **P**_*a*_ + **P**_*b*_ = 0, we get for steady state

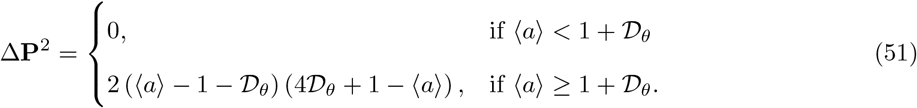

We also have, for *P≪* 1,

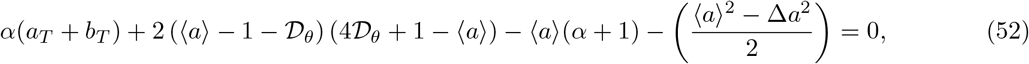

and

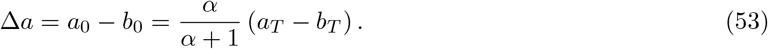

We solved these equations for ⟨*a*⟩ which showed that a small uniform increase (or decrease) in the levels of of the two proteins, while keeping that of the other one unchanged, results in a higher (lower) asymmetry in the localization of both the proteins. For large deviations from equal expression levels of two proteins, that it |*a*_*T*_ − *b*_*T*_ | ≫ 1, the nonlinear nature of *f* (*c*) will have to be taken into account.

#### 3.1.2 Non-homogeneous planar cell polarization

In the absence of expression gradients of the proteins of global module (Ft and Ds), experiments have shown a non-homogeneously polarized state of the tissue where cells become polarized but the direction of polarization is not fixed [55, 56, 5]. We consider the case of *a*_*T*_ = *b*_*T*_ = *ρ* > *ρ*_*c*_ such that the two proteins are localized asymmetrically on the cell membranes. In order to understand the nature of inhomogeneous polarization we will utilize the fact that in case of homogeneous polarization the two proteins are localized in two opposite directions. For inhomogeneous polarization we consider the case where the orientation of two proteins deviates from perfectly opposite directions by a small magnitude. That is, if *θ*_*a*_ and *θ*_*b*_ are the PCP directions for the two proteins then *θ*_*a*_(**x**) − *θ*_*b*_(**x**) = *π* + *ϵ* (**x**) where | *ϵ* | is very small. We further consider that *ρ*_0_ and magnitudes of the polarization follow *P*_*a*_ = *P*_*b*_ = *P* where *P* does not depend on location in the tissue. These assumptions makes only polarity directions to be the unknown parameters. Substitution of these assumptions in equations (31) and (32) gives us

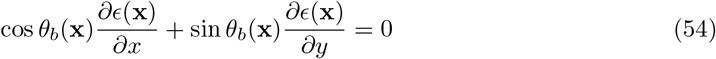

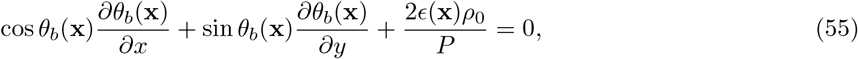

where we have kept only the leading order terms in *ϵ*. Elimination of *ϵ* from these equations give us

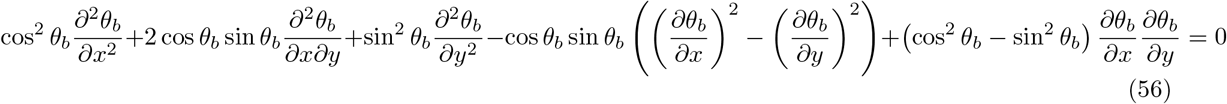

where dependency on **x** is not shown for brevity. In order to identify the nonhomogeneous polarizations it is more convenient to look at it in the polar coordinates. Transformation of above equation to polar coordinates gives

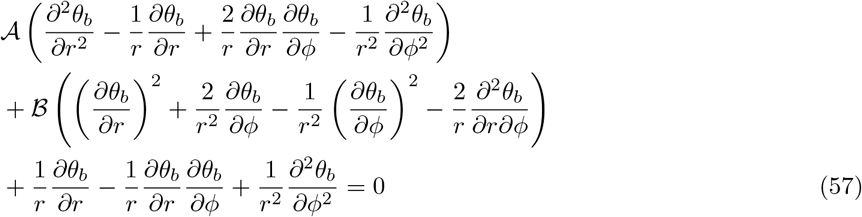

where *ϕ* is the polar angle and

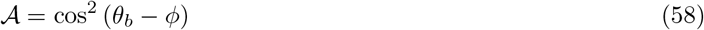

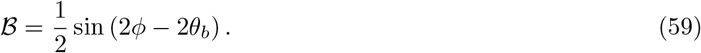

In general this equation is not analytically solvable and, therefore, we look at some special scenarios. If we focus on the region of *r* ≫ 1 and ignore the *r* dependence and consider solutions of type *θ*_*b*_(*ϕ*), the equation turns into a remarkably simpler form

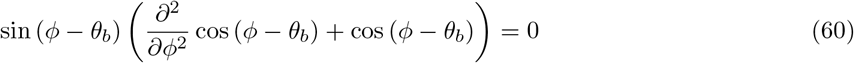

which gives us two types of solutions given by

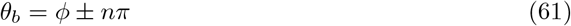

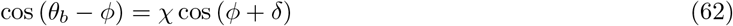

where *χ* and *δ* are arbitrary constants to be obtained from the boundary conditions. Fig. 3 shows three examples of the patterns of non-homogeneous planar cell polarity in the tissue as obtained from the analytical expressions. The numerical simulation of the discrete two dimensional model of the epithelial tissue also confirmed the existence of the non-homogeneous solutions and some such examples are shown in Fig. 3. This shows that in case of uniform protein expression the minimal continuum theory also demonstrates the non-homogeneous steady states of the PCP in epithelial tissue.

**Figure 3:**
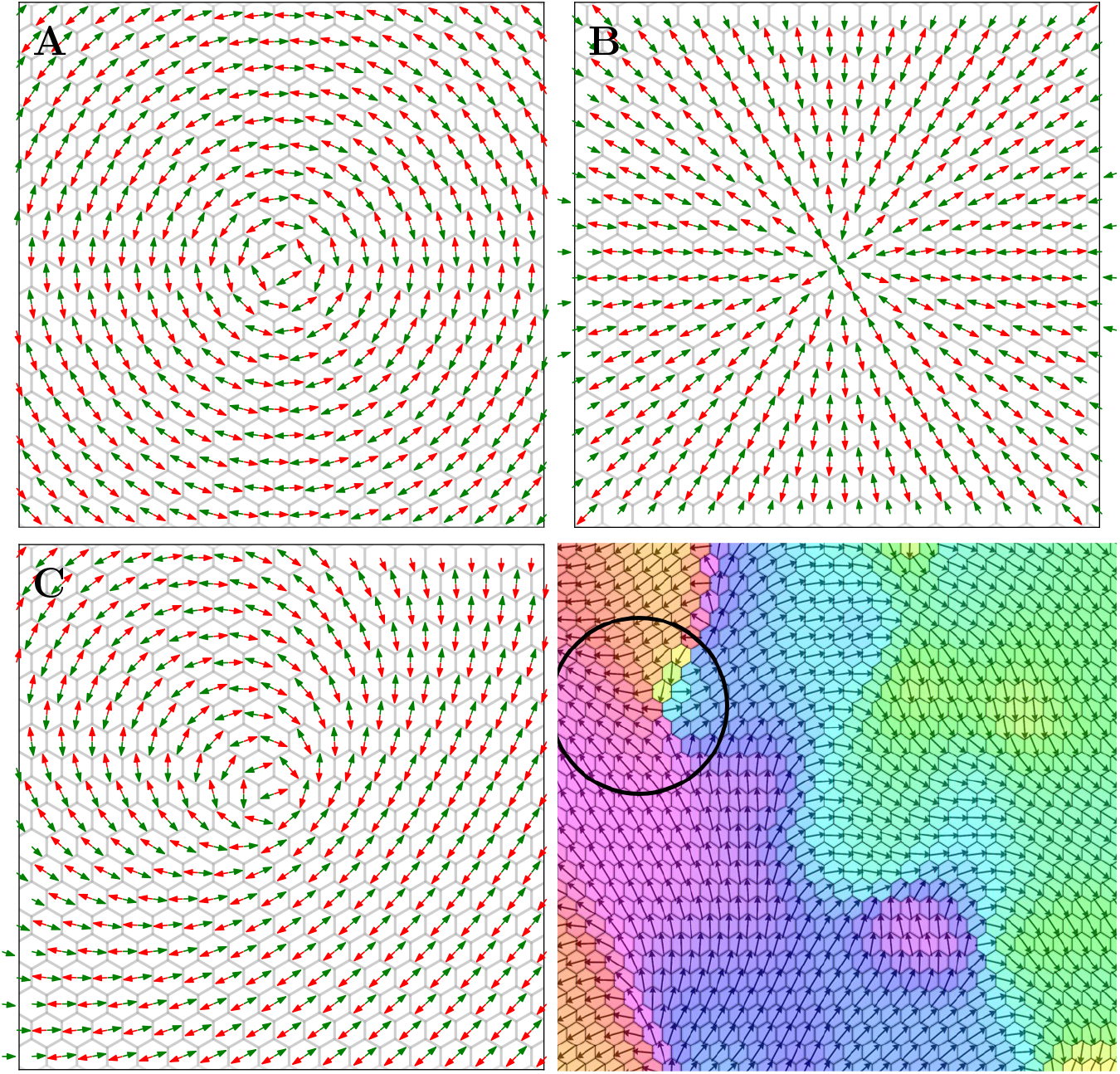
Non-homogeneous planar cell polarization with uniform protein expressions. (A-C) Three examples of non-homogeneous steady states with red and green arrows showing directions of polarization for proteins *a* and *b*, respectively, as obtained from analytical expressions. Here (A) *θ*_*b*_ = *ϕ*+*π/*2 for *χ* = 0 (swirl or vortex), (B) *θ*_*b*_ = *ϕ* + *π* (aster) and (C) *χ* = 3*/*4, *δ* = −*π/*4. (D) Non-homogeneous polarized state obtained from the numerical simulation of the discrete counterpart of the continuum model (See [51] for more details on discrete model).

### 3.2 Graded expressions of the proteins

We also consider the case where the two proteins are expressed in a tissue level gradient manner which is the case for Ft and Ds, the members of the global module [50, 18, 49]. For simplicity, we will analyze the case of graded expression of only protein *a*. The gradient is assumed to be along the *x*-axis and very shallow such that |*da/dx*| ≪ *a*_*T*_ */l*. We denote *da/dx* = *ϵ*. For this case we consider the solutions where the quantities are dependent only on *x*.

This small perturbation in the protein levels is going to affect the steady state of the system. Assuming that the shallow gradient of *a* is going to result in very small change from the homogeneous steady state, we look at the steady states with *a*_0_ = *ρ*_0_ + *a*(*x*), *b*_0_ = *ρ*_0_ + *b*(*x*) and will try to obtain the expressions for the ⟨**P**⟩ and Δ**P**. For steady state we obtain

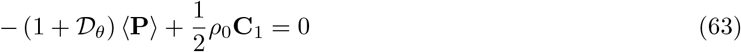

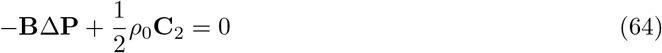

where **C**_1_ = ∇*a* + ∇*b* and **C**_2_ = ∇*b* − ∇*a*, and

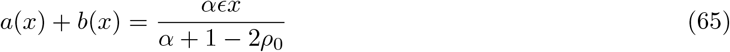

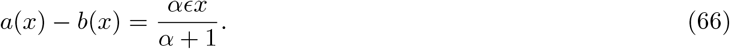

By considering ⟨**P**⟩ = ⟨*p*⟩ (cos *θ*, sin *θ*)^⊤^ and Δ**P** = Δ*p*(cos *θ*, sin *θ*)^⊤^ (with *θ* being the direction of PCP), we can see that the steady state solution corresponds to sin *θ* = 0, and we obtain

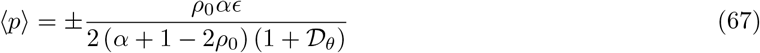

and

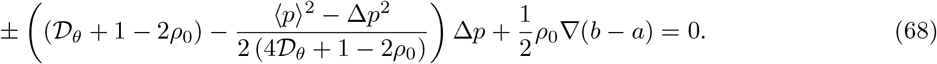

In both of above equations the prefixes ± correspond to *θ*= *π*, 0, respectively. Substituting the value of *b* − *a* gives us

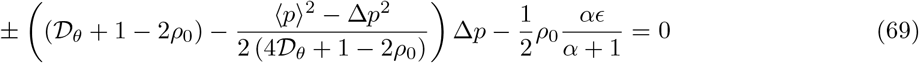

This is a cubic equation in Δ*p* with at least one real root. This implies that in the presence of expression gradient (*ϵ* ≠ 0) the unpolarized state does not exist. Furthermore, we can see that for *ρ*_0_ ≈ 0 we have 𝒟_*θ*_ + 1 − 2*ρ*_0_ > 0 and the above cubic polynomial has only one real root. However, for large *ρ*_0_, 𝒟_*θ*_ + 1 − 2*ρ*_0_ can be negative resulting is a possibility of 3 steady states. Recall that for a uniform expression levels of both proteins (that is *ϵ* = 0) it required a critical level of *ρ*_0_ for tissue polarization (albeit in arbitrary direction). Therefore, the gradient expression of protein not only polarizes the tissue in a particular direction but also facilitates the polarization itself (Fig. 4). As shown in Fig. 4A for 2*ρ*_0_ < 1 + 𝒟_*θ*_ also presence of gradient is able to polarize the tissue albeit with very small polarization magnitude *P*. For 2*ρ*_0_ > 1 + 𝒟_*θ*_, on the other hand, the cell polarization magnitude is larger for |*ϵ* | > 0 as compared to that with uniform protein expression, that is *ϵ*= 0. In both scenarios, though, the direction of the PCP is determined by the direction of the gradient (Fig. 4B). In other words, the global cue (protein expression gradient) helps PCP in its establishment and in aligning it with the tissue axis.

**Figure 4:**
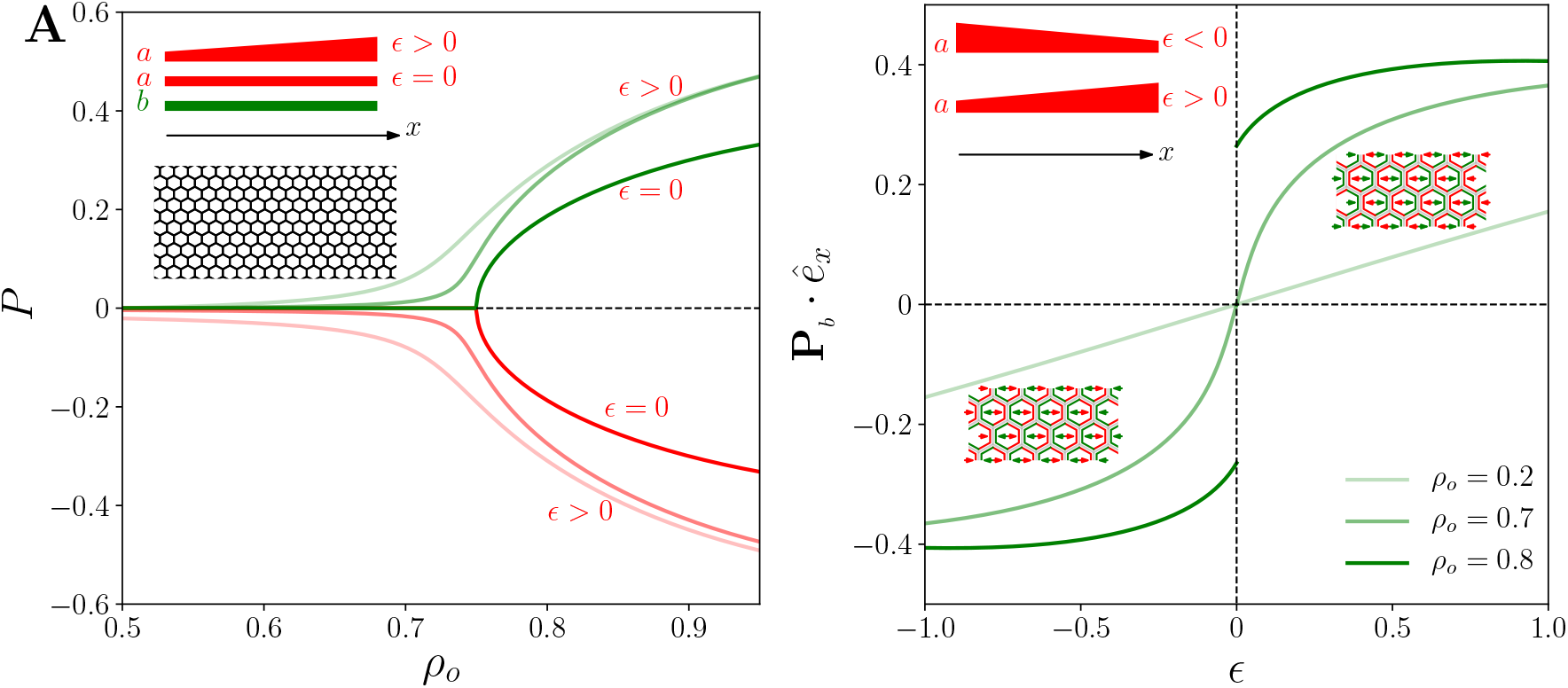
Homogeneous planar cell polarization with tissue level expression gradient of protein *a*. (A) PCP magnitude *P* as a function of *ρ*_0_ for a tissue where protein *b* is uniformly expressed but protein *a* is expressed in a tissue level gradient. Red and green curves show the asymmetry of proteins *a* and *b*, respectively. (B) Dependence of **P**_*b*_ · *ê*_*x*_ on *ϵ*, the expression gradient of *a*.

In order to assess the stability of the polarized state in the presence of the gradient expression, following the same steps as before, we obtained the dispersion relation as

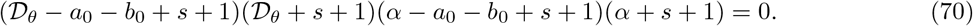

It can be seen that for *a*_0_,*b*_0 ≪_ 1 all of the roots of the above polynomial dispersion relation in *s* have negative real part. This shows that the polarized state in the presence of the protein expression gradients remains stable even against non-homogeneous perturbations. Therefore, the protein gradients also provide stability to the polarized state of the tissue.

### 3.3 Loss and overexpression of proteins in a region of the tissue

In several experimental works, the PCP proteins are removed or over-expressed in a select region (mutant clones) of the tissue to study the effect on the cells in the vicinity of that domain [36, 18, 57]. Such experimental perturbations are in contrast to the tissue wide upregulation or downregulation of the PCP proteins and uncover the local impacts of individual PCP players in polarity establishment (Fig. 5A). Using continuum theory we also looked at such localized perturbations in the epithelial tissue.

**Figure 5:**
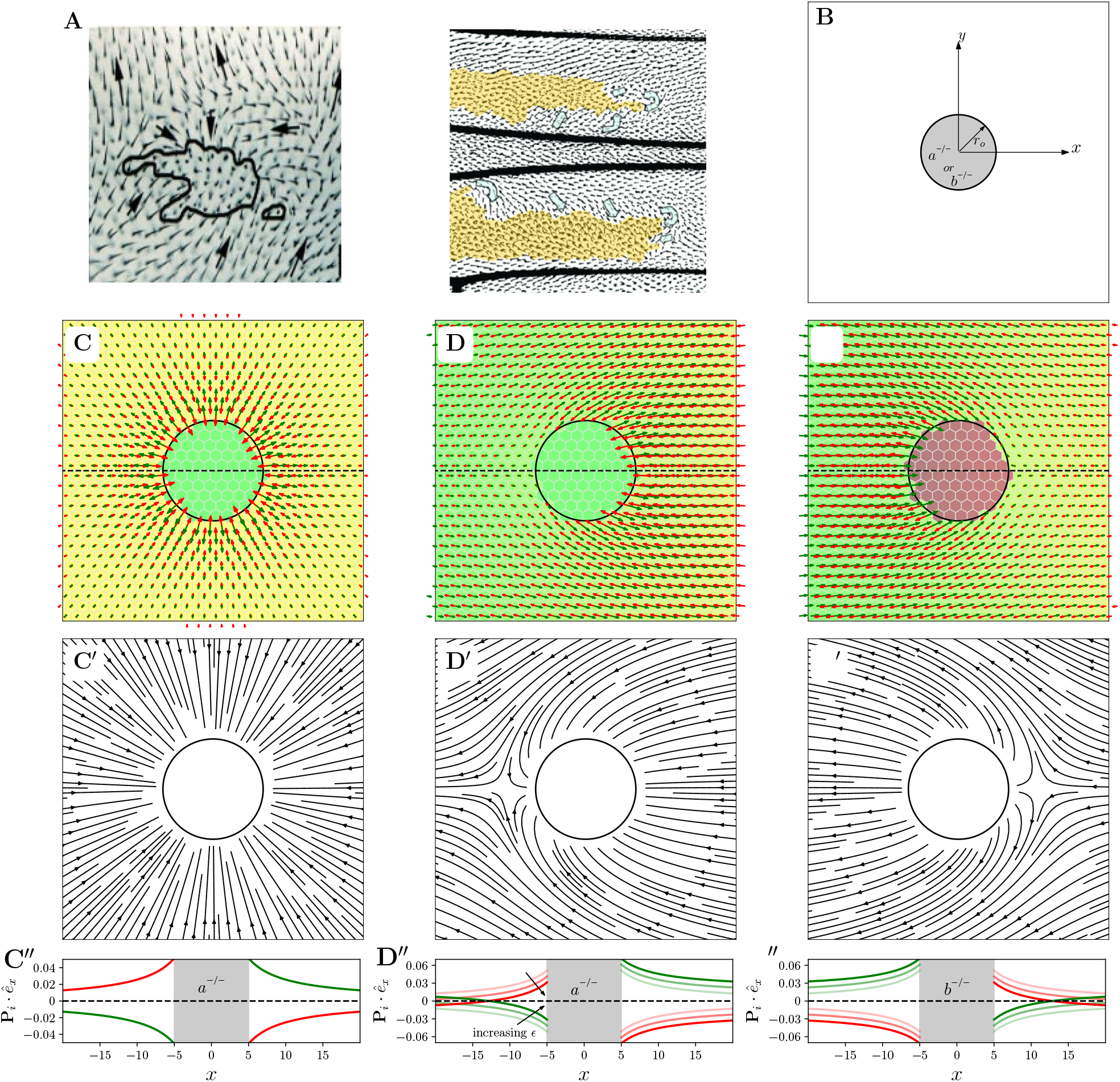
Loss of a protein from a select region of epithelial tissue. (A) Orientation of the hairs on the *Drosophila* wing when one protein in not expressed in the region of the wing. (B) Schematic representing the loss of protein *a* or *b* from the circular region (*a*^−*/*−^ or *b*^−*/*−^ clones) of epithelial tissue. Vector field showing the planar polarity of the two proteins in the tissue when (C) both *a* and *b* are expressed uniformly throughout the tissue except in the circular region where protein *a* is not present, (D) protein *b* is expressed uniformly and *a* is expressed in a linear gradient except for the circular region where *a* is not expressed, and (E) *a* and *b* are expressed in a gradient and uniform manner, respectively, except in the circular region where *b* is not expressed. (C’-E’) The streamline plots showing the orientation of the vector **P**_*a*_ − **P**_*b*_ for the cases shown in (C-E). (C”-E”) Dependence of the planar polarity (*i* = *a, b*) along the respective dashed lines marked in (C-E) on the distance from the center of the circular region.

#### 3.3.1 Loss of one protein from a circular region

We consider an idealized situation where one of the two proteins, say protein *a* for the current analysis, is selectively removed from the cells of a circular region of radius *r*_0_ in the tissue (Fig. 5B). Apart from the circular region all the other cells of the tissue have uniform distribution of both proteins. Absence of protein *a* in the cells located at the boundary of the circular domain will, by virtue of the inter-cellular protein interactions, lead to the localization of protein *b* at the boundary of circular region. In other words, the protein *b* will get radially polarized in the cells at the boundary of the circular region. This polarization will, further, polarize protein *a* in the normal cells located adjacent to the boundary of the circular domain. We further assume that the steady state solution of this system deviates from the uniform steady state solution (see Section 3.1) by a very small magnitude. We take *a*_0_ = *ρ*_0_ + *a*(*r*), *b*_0_ = *ρ*_0_ + *b*(*r*), **P**_*a*_ = **P** + *P*_*a*_(*r*)*ê*_*r*_ and **P**_*b*_ = −**P** + *P*_*b*_(*r*)*ê*_*r*_, with |**P**|, |*a*|, |*b*|, |*P*_*a*_|, |*P*_*b*_| ≪ *ρ*_0_. The assumption of the radial nature of the perturbation in the polarization is due to the fact that in the ‘normal’ cells (where both proteins are expressed in their normal levels) at the boundary of the circular region protein *a* is going to localize towards the circular region. For steady state, we substitute the aforementioned ansatz in the equations (29) (30), (31) and (32) to obtain

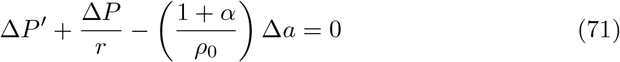

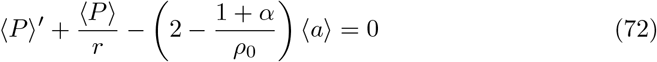

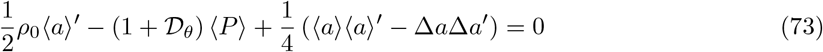

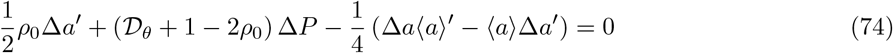

where ⟨*P*⟩ = *P*_*a*_ + *P*_*b*_, Δ*P* = *P*_*a*_ − *P*_*b*_, ⟨*a*⟩ = *a* + *b* and Δ*a* = *a* − *b*. In these equations, only the leading order terms in ⟨*P*⟩, Δ*P*, ⟨*a*⟩ and Δ*a* have been kept. Elimination of ⟨*a*⟩ from equation (72) and (73) and keeping only linear terms results in

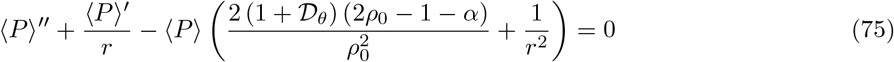

which can be rearranged to obtain following equation

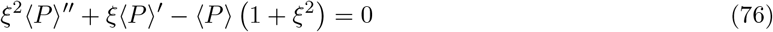

where 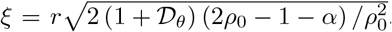. This equation is characterized as Bessel’s (for 2*ρ*_0_ < 1 + *α*) or modified Bessel’s (for 2*ρ*_0_ > 1 + *α*) equation. In order to have the polarization magnitude bounded for *r* → ∞, the solution to this equation are given by

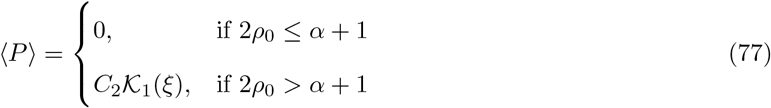

where K_1_(*ξ*) is the first order modified Bessel’s function of second kind. Similarly, from equations (71) and (74), we obtain

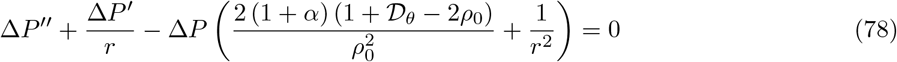

which has solution (respecting the boundary condition at *r* → ∞)

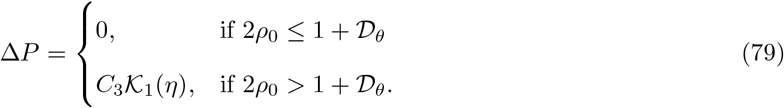

where 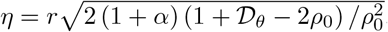. From these solutions we can obtain *P*_*a*_ = (⟨*P*⟩ + Δ*P*) */*2 and *P*_*b*_ = (⟨*P*⟩ − Δ*P*) */*2. It needs to be pointed out that the modified Bessel’s function 𝒦_1_(*r*) decays exponentially for *r* ≫ 1 which shows that similar to one-dimensional case, in two dimensions also the polarity perturbation due to localized loss of protein expression decays exponentially.

The two constants *C*_2_ and *C*_3_ are to be determined from the boundary conditions at *r* = *r*_0_. Intuitively, since protein *a* in the cells at 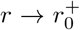 gets localized at the border of the circular region, 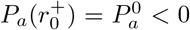. Similarly, for protein *b* we have 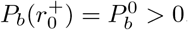. We estimate the values of 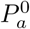 and 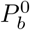 by taking advantage of the results for the case with gradient protein expressions. At the boundary of the circular region, *a* can be considered to be expressed in a gradient with gradient magnitude *ρ* since level of *a* increase from 0 to *ρ* over the length of single cell. Using equation (69), we get the values of polarization of *a* and *b* at the boundary of circular region as

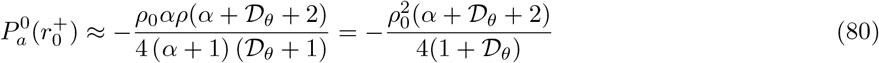

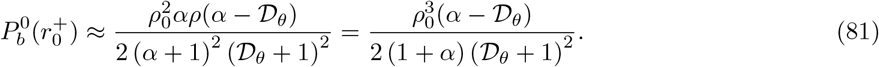

Since in the tissue with uniform protein expression the direction of polarization is arbitrary, this perturbation can results in a radial polarization near the boundary of the clone if proteins are not expressed in any gradient (Fig. 5C-C’). Although, this ‘boundary effect’ does not traverse to very large distances and decays exponentially (Fig. 5C”). Further, in the presence of a tissue level gradients in the ‘normal’ region of the tissue the effect of the loss of one protein from the clone shows different behaviors of two sides (Fig. 5D-E”). This demonstrates a domineering non-autonomy [36] or non-cell autonomous [18] effect which have been reported in experimental works on core modules of PCP [57].

#### 3.3.2 Overexpression of one protein in a circular region

Similar to the previous case of protein loss from a circular region, here also we consider the case where protein *a* is over-expressed to an amount 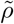 in a circular region of the tissue. In all the other cells, levels of *a* are set at a value *ρ*. The other protein *b* is uniformly expressed throughout the tissue (including the circular region) with the same value *ρ*. Similar to the protein loss case, in this scenario also, if we consider the solutions of type *a*_0_ = *ρ*_0_ + *a*(*r*), *b*_0_ = *ρ*_0_ + *b*(*r*), **P**_*a*_ = **P** + *P*_*a*_*ê*_*r*_ and **P**_*b*_ = −**P** + *P*_*b*_*ê*_*r*_ with |**P**|, |*a*|, |*b*|, |*P*_*a*_|, |*P*_*b*_| ≪ *ρ*_0_, we can see that the cells outside the circular region still follow the same equations (Equations (72), (71), (73), and (74)) as in previous case but with different boundary conditions. In this case, the polarizations of the two proteins at the boundary 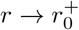 satisfy

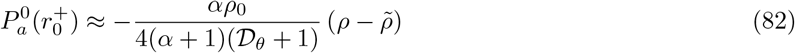

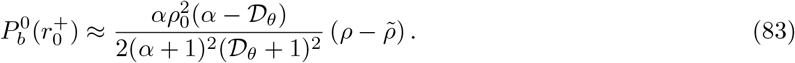

Comparing these values with those in relations (80) and (81) show that the polarization outside the circular region is opposite to that seen during protein loss from the circular region.

For the cells in the circular region, we consider solution 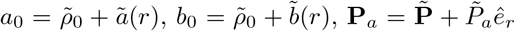 and 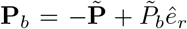 with 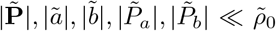. For this region also we get the similar equations as (72)-(74) with boundary conditions

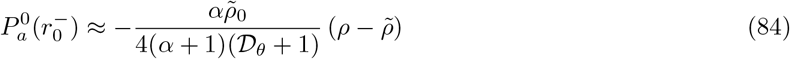

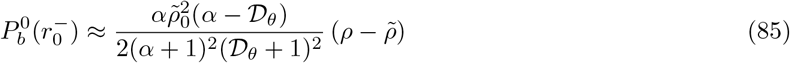

with

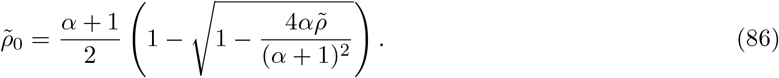

This shows that similar to the loss of one protein from a region of tissue the overexpression in a localized region also demonstrates similar effect albeit with the direction of the polarity perturbation is in the opposite direction.

Therefore, we have demonstrated that the proposed continuum model with very few parameters is able to capture most of the experimental observations. The mechanistic basis of the model formulation also helps in directly studying the experimental perturbations analytically.

## 4 Discussion

The analytical approach proposed in this work for the study of PCP presents many advantages, thanks to the consideration of the microscopic details of inter-cellular interactions, over relatively more prevalent computational discrete models [6, 36, 47, 38, 39, 40, 41]. It provides analytical understanding of the biophysics of the PCP in a planar immobile tissue and its response against perturbations similar to the genetic experiments. One can also take help of numerical methods to study the PCP in this framework in more complicated situations where sheer number of cells make numerical simulation of discrete models difficult. In addition to these advantages it also presents the PCP mechanism in a new light as discussed below.

### 4.1 Planar cell polarity as an example of active matter

The analytical continuum model of the planar cell polarity demonstrates similarities between PCP and some other extensively explored physical phenomena. Considering the cell polarization, as defined here, to be analogous to the magnetization order parameter, one can draw many parallels between PCP and classical rotor or XY model [58]. XY model is very well studied and has been known to demonstrate topological phase transitions [59, 58] when the temperature is changed. In PCP, however, we looked at the polarization mainly as a function of the concentration levels of the proteins and the inclusion of ‘temperature’ is only in the form of rotational diffusion of the polarity vector. Further, another similarity between these two systems is in terms of the external cues, that is external magnetic field in case of ferromagnetic XY model [59] and the tissue level expression gradients in case of PCP. It needs to be pointed out though that these parallels between statistical mechanics models of magnetization and PCP have also been suggested previously [37, 60, 39].

As the XY model is, in some sense, a realization of the active self propelled particles without convection [61], PCP can also be seen as the biological mechanism of the interaction among the self-propelled particles. These particles in current context are the motile epithelial cells which play crucial roles in embryonic development [62, 63, 64, 65], wound healing [65, 66, 67, 68] and cancer metastasis [65, 69, 70]. This comparison is further strengthened by our observation that the uniformly polarized state of PCP in the absence of global cue is not stable against non-homogeneous perturbations which is a well known result in hydrodynamic theory of active matter [53]. Therefore, the planar cell polarity in the epithelial tissues can be seen as another example of two dimensional active matter. This opens up several avenues for exploration and study of PCP from the active matter vantage point.

### 4.2 Inter-cellular dimer formation: ‘means to an end’ or ‘an end in itself’

Apart from the analytical framework one major deviation of the present work from the existing PCP models is the treatment of the inter-cellular protein complexes (or heterodimers) [48, 38, 71, 39, 44]. Almost all of the mathematical models of PCP in consider the cell polarity in terms of the asymmetric enrichment of the inter-cellular protein complexes or heterodimers at the cell membranes. This consideration is persistent in all previous works despite the lack of any clear experimental evidence (as far as we know) showing that it is the inter-cellular complex itself which is necessary for the smooth downstream functioning of the PCP. In contrast, we define cell polarity in terms of the asymmetric distribution of the PCP proteins on the cell periphery, irrespective of the fact if they form complex with the protein from adjacent cell or not. Instead we consider inter-cellular complex to be only a mechanism for the modulation of the binding kinetics of the membrane proteins which eventually leads to asymmetry in the protein localization on the cell membrane. This assumption is not very different from the similar considerations of intra-cellular protein-protein interactions between members of core PCP module which results in asymmetric localization of these proteins [7]. The settlement of this question of inter-cellular heterodimer being ‘means’ or ‘end’ for PCP, though, will require careful experimental effort.

### 4.3 Multiple roles of the global cue

The tissue scale gradient in the expression of Ft and Ds, the proteins from the global module of the PCP, has been shown in several developmental contexts [72, 48, 35]. The role of this gradient has been proposed to provide a global directional cue to coincide local cell polarity with the tissue axis [35]. Our analysis also attributed the role of global cue to the expression gradients but also showed that this is not its only function. As mentioned earlier, the polarized state under uniform (with no gradient) protein expressions is not stable against non-homogeneous perturbations. However, in the presence of a tissue scale expression gradient of the PCP proteins (such as Ft or Ds) the polarized state becomes stable. Therefore, expression gradients also provide stability to the polarized state in addition to being the global cue. Our results also address the question whether the global cue is necessary throughout the developmental process or only a short duration can suffice. Given the unstable nature of the polarized state under uniform protein levels the gradients are more likely to be required throughout the developmental process. A transient global cue can indeed align PCP with the tissue axis but its robustness is doubtful. This conclusion is in agreement with previous results from the discrete modeling of PCP in one dimensional model [73]. Furthermore, we also see that in the absence of the expression gradient the cell polarity (that too unstable) emerges only when the concentration of the participating proteins is above a critical level. That critical concentration, however, is not required in the presence of tissue level expression gradients for the polarity establishment. This shows that the tissue levels gradients of protein expressions play multiple roles in PCP as opposed to the conventional wisdom that its role is limited to being a global cue.

### 4.4 Limitations and future scope

In almost all developmental contexts epithelial tissues show large scales flows along with PCP and there is now ample evidence that the two processes are mechanistically linked given participation of some proteins in both processes [74]. The continuum framework presented here makes this coupling much easier by consideration of hydrodynamics on two dimensional surfaces along with PCP. Flow of the epithelial tissues is usually modeled using vertex models which are computational in nature and do not provide much analytical insight. It will be the next step in this work to look at the combined effect of PCP and flows on the overall biology and mechanics of epithelial tissues in developmental and other contexts. Furthermore, for the sake of simplicity we have also ignored many of the protein-protein interactions, especially the intra-cellular ones among the core module members such as the recruitment of the Dsh, Dgo, Vang and Pk to the respective regions of the cell boundary. These interactions can also be easily incorporated in the theory. This may, however, require keeping track of the asymmetry in multiple proteins at the cell boundaries.

Even though the continuum approach presents several advantages it also has its limitations such as the typical length scale of the features captured here is much larger than the cell size which is not always ideal from the experimentation view point. One such example is the nature of PCP near the singular point in the non-homogeneous polarization (shown in Section 3.1.2). Therefore, it will be worth while to extend this work to a hybrid approach where the small scale features can be captured using discrete model and large scale epithelial biomechanics can be explored using continuum framework.

Further, it has to be noted that few PCP proteins play other non-PCP related functions as well which may influence some of the model assumptions if they are removed from a region of the tissue. For an example, Ft, a tumor suppressor protein which has also been shown to affect the cell shapes and sizes in epithelial tissue [75]. Therefore, the modeling of any perturbation in the levels of such proteins in select regions of the tissue should ideally be accompanied by incorporating such geometrical factors as well which is not considered here.

## Acknowledgments

We would like to acknowledge and thank Prof Sriram Ramaswamy for his constant valuable support and inputs. D.S. is supported by KVPY fellowship awarded by Department of Science and Technology (DST), Government of India. MKJ is supported by Ramanujan Fellowship (SB/S2/RJN 049/2018) awarded by the Science and Engineering Research Board (SERB), DST, Government of India. MSR would like to acknowledge the financial support provided by the seed grant from IIT Hyderabad.

## References

[1] U. Tepass, G. Tanentzapf, R. Ward, and R. Fehon. Epithelial cell polarity and cell junctions in Drosophila. Annu Rev Genet, 35:747–784, 2001.

[2] M. C. Gibson and N. Perrimon. Apicobasal polarization: epithelial form and function. Curr Opin Cell Biol, 15(6):747–752, Dec 2003.

[3] M. Simons and M. Mlodzik. Planar cell polarity signaling: from fly development to human disease. Annu Rev Genet, 42:517–540, 2008.

[4] M. T. Butler and J. B. Wallingford. Planar cell polarity in development and disease. Nat Rev Mol Cell Biol, 18(6):375–388, 06 2017.

[5] D. Devenport. The cell biology of planar cell polarity. J Cell Biol, 207(2):171–179, Oct 2014.

[6] J. D. Axelrod and C. J. Tomlin. Modeling the control of planar cell polarity. Wiley Interdiscip Rev Syst Biol Med, 3(5):588–605, 2011.

[7] Ying Peng and Jeffrey D. Axelrod. Chapter two - asymmetric protein localization in planar cell polarity: Mechanisms, puzzles, and challenges. In Yingzi Yang, editor, Planar Cell Polarity During Development, volume 101 of Current Topics in Developmental Biology, pages 33–53. Academic Press, 2012.

[8] A. Brittle, C. Thomas, and D. Strutt. Planar polarity specification through asymmetric subcellular localization of Fat and Dachsous. Curr Biol, 22(10):907–914, May 2012.

[9] Z. Kibar, K. J. Vogan, N. Groulx, M. J. Justice, D. A. Underhill, and P. Gros. Ltap, a mammalian homolog of Drosophila Strabismus/Van Gogh, is altered in the mouse neural tube mutant Loop-tail. Nat Genet, 28(3):251–255, Jul 2001.

[10] J. N. Murdoch, K. Doudney, C. Paternotte, A. J. Copp, and P. Stanier. Severe neural tube defects in the loop-tail mouse result from mutation of Lpp1, a novel gene involved in floor plate specification. Hum Mol Genet, 10(22):2593–2601, Oct 2001.

[11] J. A. Curtin, E. Quint, V. Tsipouri, R. M. Arkell, B. Cattanach, A. J. Copp, D. J. Henderson, N. Spurr, P. Stanier, E. M. Fisher, P. M. Nolan, K. P. Steel, S. D. Brown, I. C. Gray, and J. N. Murdoch. Mutation of Celsr1 disrupts planar polarity of inner ear hair cells and causes severe neural tube defects in the mouse. Curr Biol, 13(13):1129–1133, Jul 2003.

[12] S. K. Kim, A. Shindo, T. J. Park, E. C. Oh, S. Ghosh, R. S. Gray, R. A. Lewis, C. A. Johnson, T. Attie-Bittach, N. Katsanis, and J. B. Wallingford. Planar cell polarity acts through septins to control collective cell movement and ciliogenesis. Science, 329(5997):1337–1340, Sep 2010.

[13] H. Song, J. Hu, W. Chen, G. Elliott, P. Andre, B. Gao, and Y. Yang. Planar cell polarity breaks bilateral symmetry by controlling ciliary positioning. Nature, 466(7304):378–382, Jul 2010.

[14] E. K. Vladar, R. D. Bayly, A. M. Sangoram, M. P. Scott, and J. D. Axelrod. Microtubules enable the planar cell polarity of airway cilia. Curr Biol, 22(23):2203–2212, Dec 2012.

[15] Y. Mao, A. L. Tournier, P. A. Bates, J. E. Gale, N. Tapon, and B. J. Thompson. Planar polarization of the atypical myosin Dachs orients cell divisions in Drosophila. Genes Dev, 25(2):131–136, Jan 2011.

[16] D. Devenport and E. Fuchs. Planar polarization in embryonic epidermis orchestrates global asymmetric morphogenesis of hair follicles. Nat Cell Biol, 10(11):1257–1268, Nov 2008.

[17] F. Mangione and E. Mart?n-Blanco. The Dachsous/Fat/Four-Jointed Pathway Directs the Uniform Axial Orientation of Epithelial Cells in the Drosophila Abdomen. Cell Rep, 25(10):2836–2850, 12 2018.

[18] Maja Matis and J. D. Axelrod. Regulation of pcp by the fat signaling pathway. Genes Dev, 27(20):2207–2220, 10 2013.

[19] Y. Yang and M. Mlodzik. Wnt-Frizzled/planar cell polarity signaling: cellular orientation by facing the wind (Wnt). Annu Rev Cell Dev Biol, 31:623–646, 2015.

[20] T. Usui, Y. Shima, Y. Shimada, S. Hirano, R. W. Burgess, T. L. Schwarz, M. Takeichi, and T. Uemura. Flamingo, a seven-pass transmembrane cadherin, regulates planar cell polarity under the control of Frizzled. Cell, 98(5):585–595, Sep 1999.

[21] C. J. Formstone and I. Mason. Combinatorial activity of Flamingo proteins directs convergence and extension within the early zebrafish embryo via the planar cell polarity pathway. Dev Biol, 282(2):320–335, Jun 2005.

[22] J. M. Carvajal-Gonzalez and M. Mlodzik. Mechanisms of planar cell polarity establishment in Drosophila. F1000Prime Rep, 6:98, 2014.

[23] M. T. Butler and J. B. Wallingford. Planar cell polarity in development and disease. Nat. Rev. Mol. Cell Biol., 18(6):375–388, 06 2017.

[24] J. Wu and M. Mlodzik. The frizzled extracellular domain is a ligand for Van Gogh/Stbm during nonautonomous planar cell polarity signaling. Dev Cell, 15(3):462–469, Sep 2008.

[25] D. R. Tree, J. M. Shulman, R. Rousset, M. P. Scott, D. Gubb, and J. D. Axelrod. Prickle mediates feedback amplification to generate asymmetric planar cell polarity signaling. Cell, 109(3):371–381, May 2002.

[26] H. Strutt and D. Strutt. Differential stability of flamingo protein complexes underlies the establishment of planar polarity. Curr Biol, 18(20):1555–1564, Oct 2008.

[27] G. Mottola, A. K. Classen, M. Gonz?lez-Gait?n, S. Eaton, and M. Zerial. A novel function for the Rab5 effector Rabenosyn-5 in planar cell polarity. Development, 137(14):2353–2364, Jul 2010.

[28] P. A. Lawrence, J. Casal, and G. Struhl. Cell interactions and planar polarity in the abdominal epidermis of Drosophila. Development, 131(19):4651–4664, Oct 2004.

[29] R. Bastock, H. Strutt, and D. Strutt. Strabismus is asymmetrically localised and binds to Prickle and Dishevelled during Drosophila planar polarity patterning. Development, 130(13):3007–3014, Jul 2003.

[30] H. Strutt, S. J. Warrington, and D. Strutt. Dynamics of core planar polarity protein turnover and stable assembly into discrete membrane subdomains. Dev Cell, 20(4):511–525, Apr 2011.

[31] D. Ma, C. H. Yang, H. McNeill, M. A. Simon, and J. D. Axelrod. Fidelity in planar cell polarity signalling. Nature, 421(6922):543–547, Jan 2003.

[32] M. Matis, D. A. Russler-Germain, Q. Hu, C. J. Tomlin, and J. D. Axelrod. Microtubules provide directional information for core PCP function. Elife, 3:e02893, Aug 2014.

[33] T. Ayukawa, M. Akiyama, J. L. Mummery-Widmer, T. Stoeger, J. Sasaki, J. A. Knoblich, H. Senoo, T. Sasaki, and M. Yamazaki. Dachsous-dependent asymmetric localization of spiny-legs determines planar cell polarity orientation in Drosophila. Cell Rep, 8(2):610–621, Jul 2014.

[34] J. Casal, P. A. Lawrence, and G. Struhl. Two separate molecular systems, Dachsous/Fat and Starry night/Frizzled, act independently to confer planar cell polarity. Development, 133(22):4561–4572, Nov 2006.

[35] H. Matakatsu and S. S. Blair. Interactions between Fat and Dachsous and the regulation of planar cell polarity in the Drosophila wing. Development, 131(15):3785–3794, Aug 2004.

[36] K. Amonlirdviman, N. A. Khare, D. R. P. Tree, W.-S. Chen, J. D. Axelrod, and C. J. Tomlin. Mathematical modeling of planar cell polarity to understand domineering nonautonomy. Science, 307(5708):423–426, 2005.

[37] Y. Burak and B. I. Shraiman. Order and stochastic dynamics in Drosophila planar cell polarity. PLoS Comput. Biol., 5(12):e1000628, Dec 2009.

[38] M. Mani, S. Goyal, K. D. Irvine, and B. I. Shraiman. Collective polarization model for gradient sensing via Dachsous-Fat intercellular signaling. Proc. Natl. Acad. Sci. U.S.A., 110(51):20420–20425, Dec 2013.

[39] L. D. Hazelwood and J. M. Hancock. Functional modelling of planar cell polarity: an approach for identifying molecular function. BMC Dev. Biol., 13:20, May 2013.

[40] Mohit Kumar Jolly, Mohd Suhail Rizvi, Amit Kumar, and Pradip Sinha. Mathematical modeling of sub-cellular asymmetry of fat-dachsous heterodimer for generation of planar cell polarity. PLOS ONE, 9(5):1–10, 05 2014.

[41] K. H. Fisher and D. Strutt. A theoretical framework for planar polarity establishment through interpretation of graded cues by molecular bridges. Development, 146(3), 02 2019.

[42] Dali Ma, Keith Amonlirdviman, Robin L. Raffard, Alessandro Abate, Claire J. Tomlin, and Jeffrey D. Axelrod. Cell packing influences planar cell polarity signaling. Proceedings of the National Academy of Sciences, 105(48):18800–18805, 2008.

[43] Guillaume Salbreux, Linda K. Barthel, Pamela A. Raymond, and David K. Lubensky. Coupling mechanical deformations and planar cell polarity to create regular patterns in the zebrafish retina. PLOS Computational Biology, 8(8):1–20, 08 2012.

[44] Shahriar Shadkhoo and Madhav Mani. The role of intracellular interactions in the collective polarization of tissues and its interplay with cellular geometry. PLOS Computational Biology, 15(11):1–29, 11 2019.

[45] Y. Shimada, S. Yonemura, H. Ohkura, D. Strutt, and T. Uemura. Polarized transport of Frizzled along the planar microtubule arrays in Drosophila wing epithelium. Dev Cell, 10(2):209–222, Feb 2006.

[46] T. Harumoto, M. Ito, Y. Shimada, T. J. Kobayashi, H. R. Ueda, B. Lu, and T. Uemura. Atypical cadherins Dachsous and Fat control dynamics of noncentrosomal microtubules in planar cell polarity. Dev Cell, 19(3):389–401, Sep 2010.

[47] Y. Burak and B. I. Shraiman. Order and stochastic dynamics in Drosophila planar cell polarity. PLoS Comput Biol, 5(12):e1000628, Dec 2009.

[48] R. Hale, A. L. Brittle, K. H. Fisher, N. A. Monk, and D. Strutt. Cellular interpretation of the long-range gradient of Four-jointed activity in the Drosophila wing. Elife, 4, Feb 2015.

[49] G. M. Collu and M. Mlodzik. Planar polarity: converting a morphogen gradient into cellular polarity. Curr Biol, 25(9):R372–374, May 2015.

[50] P. A. Lawrence, G. Struhl, and J. Casal. Do the protocadherins Fat and Dachsous link up to determine both planar cell polarity and the dimensions of organs? Nat Cell Biol, 10(12):1379–1382, Dec 2008.

[51] D. Singh, S. Ramaswamy, M. K. Jolly, and M. S. Rizvi. Emergent dynamics of the interplay of local and global interactions in planar cell polarity. Submitted.

[52] Justin Hogan, Meagan Valentine, Chris Cox, Kristy Doyle, and Simon Collier. Two frizzled planar cell polarity signals in the drosophila wing are differentially organized by the fat/dachsous pathway. PLOS Genetics, 7(2):1–14, 02 2011.

[53] E. Bertin, M. Droz, and G. Gr?goire. Boltzmann and hydrodynamic description for self-propelled particles. Phys Rev E Stat Nonlin Soft Matter Phys, 74(2 Pt 1):022101, Aug 2006.

[54] Eric Bertin, Hugues Chaté, Francesco Ginelli, Shradha Mishra, Anton Peshkov, and Sriram Ramaswamy. Mesoscopic theory for fluctuating active nematics. New Journal of Physics, 15(8):085032, aug 2013.

[55] Y. Wang, H. Chang, and J. Nathans. When whorls collide: the development of hair patterns in frizzled 6 mutant mice. Development, 137(23):4091–4099, Dec 2010.

[56] M. Cetera, L. Leybova, F. W. Woo, M. Deans, and D. Devenport. Planar cell polarity-dependent and independent functions in the emergence of tissue-scale hair follicle patterns. Dev Biol, 428(1):188–203, 08 2017.

[57] M. L. Stoller, O. Roman, and M. R. Deans. Domineering non-autonomy in Vangl1;Vangl2 double mutants demonstrates intercellular PCP signaling in the vertebrate inner ear. Dev Biol, 437(1):17–26, May 2018.

[58] Moshe Gitterman. Phase Transitions: Modern Applications. World Scientific, 2 edition.

[59] M E Gouvea, F G Mertens, A R Bishop, and G M Wysin. The classical two-dimensional XY model with in-plane magnetic field. 2(7):1853–1868, feb 1990.

[60] K. B. Hoffmann, A. Voss-Böhme, J. C. Rink, and L. Brusch. A dynamically diluted alignment model reveals the impact of cell turnover on the plasticity of tissue polarity patterns. J R Soc Interface, 14(135), 10 2017.

[61] András Czirók, H Eugene Stanley, and Tamás Vicsek. Spontaneously ordered motion of self-propelled particles. 30(5):1375–1385, mar 1997.

[62] R. G. Morris and M. Rao. Active morphogenesis of epithelial monolayers. Phys Rev E, 100(2-1):022413, Aug 2019.

[63] L. A. Davidson. Epithelial machines that shape the embryo. Trends Cell Biol, 22(2):82–87, Feb 2012.

[64] David Shook and Ray Keller. Mechanisms, mechanics and function of epithelial–mesenchymal transitions in early development. Mechanisms of Development, 120(11):1351–1383, 2003. The Cell in Development.

[65] D. H. Kim, T. Xing, Z. Yang, R. Dudek, Q. Lu, and Y. H. Chen. Epithelial Mesenchymal Transition in Embryonic Development, Tissue Repair and Cancer: A Comprehensive Overview. J Clin Med, 7(1), Dec 2017.

[66] A. Brugués, E. Anon, V. Conte, J. H. Veldhuis, M. Gupta, J. Colombelli, J. J. Muñoz, G. W. Brodland, Ladoux, and X. Trepat. Forces driving epithelial wound healing. Nat Phys, 10(9):683–690, Sep 2014.

[67] N. D. Evans, R. O. Oreffo, E. Healy, P. J. Thurner, and Y. H. Man. Epithelial mechanobiology, skin wound healing, and the stem cell niche. J Mech Behav Biomed Mater, 28:397–409, Dec 2013.

[68] G. Leoni, P. A. Neumann, R. Sumagin, T. L. Denning, and A. Nusrat. Wound repair: role of immune-epithelial interactions. Mucosal Immunol, 8(5):959–968, Sep 2015.

[69] J. M. Northcott, I. S. Dean, J. K. Mouw, and V. M. Weaver. Feeling Stress: The Mechanics of Cancer Progression and Aggression. Front Cell Dev Biol, 6:17, 2018.

[70] D. Ribatti, R. Tamma, and T. Annese. Epithelial-Mesenchymal Transition in Cancer: A Historical Overview. Transl Oncol, 13(6):100773, Jun 2020.

[71] K. Amonlirdviman, N. A. Khare, D. R. Tree, W. S. Chen, J. D. Axelrod, and C. J. Tomlin. Mathematical modeling of planar cell polarity to understand domineering nonautonomy. Science, 307(5708):423–426, Jan 2005.

[72] A. Kumar, M. S. Rizvi, T. Athilingam, S. S. Parihar, and P. Sinha. thorax morphogenesis. Mol Biol Cell, 31(7):546–560, 03 2020.

[73] S. Fischer, P. Houston, N. A. Monk, and M. R. Owen. Is a persistent global bias necessary for the establishment of planar cell polarity? PLoS One, 8(4):e60064, 2013.

[74] B. Aigouy, R. Farhadifar, D. B. Staple, A. Sagner, J. C. Röper, F. Jülicher, and S. Eaton. Cell flow reorients the axis of planar polarity in the wing epithelium of Drosophila. Cell, 142(5):773–786, Sep 2010.

[75] M. Jaiswal, N. Agrawal, and P. Sinha. Fat and Wingless signaling oppositely regulate epithelial cell-cell adhesion and distal wing development in Drosophila. Development, 133(5):925–935, Mar 2006.

